# LPS/TLR4-Activating Exosomes from M1-Polarized Avian Macrophages Enhances IBV Vaccine-Induced Protective Immunity in Chickens

**DOI:** 10.1101/2025.04.08.647822

**Authors:** Jun Zhou, Shikai Cai, Hongbin Huang, Fan Yang, Kexin Pan, Zhaoyang Sun, Yanfeng Chen, Yun Fan, Feng Wen, Limei Qin, Yun Zhang

**Affiliations:** Guangdong Provincial Key Laboratory of Animal Molecular Design and Precise Breeding, School of Animal Science and Technology, Foshan University, Foshan, China; School of Biomedical Engineering, Hainan University, Haikou, China

**Keywords:** Avian Infectious Bronchitis, Macrophage, Exosomes, Immunostimulation, Adjuvant, Attenuated Vaccine

## Abstract

Infectious bronchitis virus (IBV) presents a substantial economic burden to poultry production due to extensive serotypic diversity and limited cross-protection afforded by conventional vaccines. This study evaluated exosomes derived from M1-polarized chicken macrophages (HD11_M1_-exo) as a novel adjuvant for IBV vaccination. HD11_M1_-exo, isolated from LPS-activated HD11 macrophages via ultracentrifugation, demonstrated significant immunomodulatory properties across multiple experimental systems. In vitro analyses revealed HD11_M1_-exo enhanced macrophage phagocytosis and cellular immune activation via the LPS/TLR4 signaling pathway. In embryonated eggs, HD11_M1_-exo pre-treatment induced TNF-α upregulation, enhanced viral resistance, and reduced pathological damage. In chickens, HD11_M1_-exo administration elevated CD80/CD86 and TGF-β4 expression in respiratory tissues and increased secretory IgA in lacrimal fluid. When combined with H120 vaccine, HD11_M1_-exo significantly augmented both humoral immunity (elevated serum IgY and mucosal IgA) and cellular responses (increased CD80/CD86 expression), surpassing commercial adjuvants in efficacy. Following viral challenge, HD11_M1_-exo+H120 immunized chickens exhibited significantly reduced viral loads and attenuated histopathology compared to controls. These data collectively suggest that exosome-based formulations may serve as effective adjuvants for enhancing poultry vaccine immunogenicity and protective efficacy.

## INTRODUCTION

Avian infectious bronchitis (**IB**), an acute and highly contagious disease caused by infectious bronchitis virus (**IBV**), is characterized by respiratory tract lesions, nephritis, and reproductive tract abnormalities resulting in reduced egg production in layers, manifesting as a complex syndrome with substantial economic consequences for the global poultry industry (Bande et al., 2017). IBV is an enveloped, pleomorphic virus belonging to the genus Gammacoronavirus within the family Coronaviridae, possessing a linear, single-stranded, positive-sense RNA genome of approximately 27.5-28 kb in length (Falchieri et al., 2024). The remarkable genetic plasticity of IBV facilitates ongoing evolutionary adaptation through both point mutations in the spike glycoprotein and homologous recombination events, resulting in the persistent emergence of novel genetic variants and epidemiologically distinct serotypes in the field. Moreover, the serotype-specific immunity elicited by a single IBV serotype confers limited heterologous protection against antigenically diverse field strains (Zhao et al., 2023). This immunological constraint poses substantial challenges for the development of broadly protective vaccines, complicating effective control strategies against IBV with its considerable antigenic and epidemiological heterogeneity.

Exosomes are small cell-derived extracellular vesicles originating from multivesicular bodies, ubiquitous in biological fluids including plasma, lymph, saliva, semen, urine, and cerebrospinal fluid (Bakhshandeh et al., 2017; Lu et al., 2018) These membranous nanovesicles typically measure 40∼160 nm in diameter with a characteristic buoyant density of 1.13∼1.19 g/mL under ultracentrifugation conditions. Transmission electron microscopy demonstrates their distinctive cup-shaped or spherical morphology with lipid bilayer boundaries (Deb et al., 2021). Exosomes carry a multifaceted molecular payload from progenitor cells, including functional proteins, bioactive lipids, and regulatory miRNAs, which are selectively packaged and can be transferred to recipient cells through membrane fusion or endocytic uptake mechanisms. These biologically active nanovesicles serve as critical mediators in numerous physiological and pathological processes by orchestrating intercellular signaling networks, regulating immune system activation and suppression, directing cellular motility and tissue infiltration, facilitating neovascularization, and modulating pathogenic processes in various disease conditions (Mathivanan et al., 2010; Gross et al., 2012). Macrophages, serving as ubiquitous sentinel cells throughout tissues, constitute essential components of tissue homeostasis and host defense mechanisms against pathogen invasion (Davies et al., 2013). Tissue-resident macrophages exhibit remarkable plasticity and polarize into functionally distinct phenotypes in response to microenvironmental cues, conventionally categorized as classically activated (M1) macrophages driving pro-inflammatory responses and alternatively activated (M2) macrophages associated with immunoregulation and tissue repair (Martinez and Gordon, 2014). Experimental studies have established that macrophage polarization is triggered by specific molecular stimuli: lipopolysaccharide (**LPS**) induces M1 phenotype development, whereas interleukin-4 (IL-4) directs M2 phenotype differentiation (Sica and Mantovani, 2012). During immune defense responses, M1 macrophages demonstrate bifunctional effector mechanisms. These cells exert direct antimicrobial activity through the generation of reactive oxygen species (**ROS**) that compromise microbial integrity and through pathogen phagocytosis. Concurrently, they mediate indirect immunoregulatory functions via cytokine secretion and associated signaling pathways (Patel et al., 2017).

Exosomes secreted by M1 macrophages transport bioactive molecules from progenitor cells mediating immunostimulatory activities. Hong et al. showed that dsRNA virus-infected chicken macrophages activated via TLR3 ligands coordinate innate immune responses in naïve macrophages and T lymphocytes through NF-κB signaling cascade modulation (Hong et al., 2021b). Furthermore, Hong et al. systematically characterized the functional capacity of macrophage-derived exosomes in avian immune responses at the cellular level. Their findings documented that LPS-stimulated exosomes possess minimal immunogenicity, increased cellular uptake, and immune-modulating characteristics, promoting immune activation and cytokine secretion through MyD88-dependent NF-κB signaling, thus establishing them as potent immunostimulatory agents (Hong et al., 2021a). Srinivasan et al. elucidated that exosomes secreted by poly(I:C)-activated macrophages drive M1 phenotype differentiation in murine lymph node-resident macrophages and trigger NF-κB signaling cascade activation (Srinivasan et al., 2017). Tang et al. documented that exosomes secreted by LPS-activated human monocytes upregulate CCL2, ICAM-1, and IL-6 expression in endothelial cells mediated via the NF-κB signaling cascade (Tang et al., 2016). Collectively, exosomes derived from LPS-polarized M1 macrophages demonstrate minimal intrinsic immunogenicity, enhanced cellular internalization, and potent immunomodulatory properties. Through activation of specific intracellular signaling cascades, these vesicles augment immune responses and cytokine secretion. These biological nanoparticles represent promising vaccine adjuvant candidates with target specificity, precision delivery capability, and immunostimulatory efficacy. In the present investigation, LPS stimulation was utilized to induce M1 phenotype differentiation in avian HD11 macrophages, with subsequent exosome purification from culture supernatants through differential ultracentrifugation. We developed a comprehensive evaluation framework integrating in vitro cellular, embryonated egg, and in vivo avian models to systematically evaluate the immunostimulatory properties and mechanistic basis of M1-polarized avian macrophage-derived exosomes (**HD11_M1_-exo**) and quantify their adjuvant synergy with attenuated IB vaccine, benchmarked against commercial immunopotentiators.

## Materials and Methods

### Cells, chicken embryos, and animals

The HD11 cell line was maintained by the Guangdong Provincial Key Laboratory of Animal Molecular Design and Precise Breeding, Foshan University. One-day-old chicks and fertilized chicken embryos were purchased from Nanhai Poultry Breeding Co., Ltd., Foshan City, Guangdong Province.

### Viruses and reagents

The IBV attenuated vaccine (H120 strain) and the IBV-HX strain (GenBank accession number: OP846990) are preserved by the Guangdong Provincial Key Laboratory of Animal Molecular Design and Precision Breeding at Foshan University. Lipopolysaccharide (LPS) was purchased from Shanghai Biyuntian Biotechnology Co., Ltd. (Shanghai, China); Chicken Immunoglobulin A (IgA) ELISA kit was obtained from Quanzhou Ruixin Biotechnology Co., Ltd. (Fujian, China); Aluminum hydroxide adjuvant was acquired from Beijing Boao Long Biotechnology Co., Ltd. (Beijing, China); Commercial avian adjuvant (IMS 1313 VG N ST, referred to as 1313) was sourced from SEPPIC Special Chemicals Co., Ltd. (Shanghai, China); Chicken Infectious Bronchitis Antibody Test Kit (Indirect ELISA) was purchased from Shenzhen Zhenrui Biotechnology Co., Ltd. (Shenzhen, China).

### Exosome isolation and identification

To obtain exosomes from HD11_M1_-exo, when the density of HD11 cells reaches 80% and they are in good growth condition, wash the cells three times with PBS to remove the original culture medium, then replace it with RPMI 1640 complete medium containing 1% exosome-depleted serum. Randomly select one bottle of cells as a blank control group, and add LPS to the remaining culture bottles to a final concentration of 1 μg/mL. Incubate all culture bottles at 37 °C with 5% CO2 for 24 hours. For the isolation of HD11_M1_-derived exosomes, avian macrophage HD11 cells were maintained until reaching 80% confluence and exhibiting log-phase growth. Cultures were rinsed thrice with sterile PBS (pH 7.4) to eliminate residual growth medium, followed by supplementation with RPMI 1640 medium containing 1% exosome-depleted FBS. Cell cultures were designated as either non-stimulated controls or experimental groups that received LPS stimulation (1 μg/mL). All cultures were subsequently incubated at 37°C in a humidified atmosphere containing 5% CO₂ for 24 h. After incubation, collect the HD11 cells from both the control group and the treated group, as well as the supernatant from the treated group cells. Use qPCR to identify typical markers of M1 macrophages (INOS, IL-6, IL-12, IL-23, CD80, CD86) to confirm macrophage polarization to the M1 type. Then, use ultracentrifugation to isolate exosomes from the cell supernatant, and conduct transmission electron microscopy, particle size analysis, and Western Blot experiments to characterize the extracted exosomes.

### Cellular Experiment Design

Avian macrophage HD11 cells were seeded in 96-well microplates (1×10⁵ cells/well) and maintained until reaching 80% confluence under standard culture conditions. Cells were subsequently assigned to either negative control (NC) or exosome-treated (Exo) groups, with triplicate wells per group. The Exo group received 20 μg/mL HD11_M1_-derived exosomes in serum-free medium, while the NC group was administered an equivalent volume of sterile PBS (pH 7.4). Following 24 h incubation at 37°C in a humidified 5% CO₂ atmosphere, culture supernatants were aspirated. For the neutral red uptake assay, a 0.1% (w/v) neutral red solution was freshly prepared by dissolving neutral red powder in sterile DPBS and filtering through a 0.22 μm membrane. Add 100 μL to each well to achieve a final concentration of neutral red at 1 μg/mL in each well. Incubate the culture plates in a 37 ℃ incubator for an additional 1 hour. Subsequently, wash the wells three times with PBS and add 100 μL of cell lysis buffer (a mixture of ice acetic acid and anhydrous ethanol in a 1:1 ratio) to each well. Gently shake for 15 seconds and let stand at room temperature for 2 hours to allow complete cell lysis. Finally, measure the absorbance of each well at a wavelength of 540 nm.

HD11 cells in good growth condition were seeded into a 24-well culture plate. The experiment was divided into a negative control group (NC group) and an exosome treatment group (Exo group), with three independent repeats for each group. When the cell confluence reached approximately 80%, the Exo group was stimulated with 20 μg/mL of HD11_M1_-exo for 24 hours, while the NC group was treated with an equal volume of PBS. The transcriptional levels of the LPS/TLR4 signaling pathway-related genes CD14, TLR4, TRAM, TRAF3, IRF3, and CD80 were detected by qPCR.

### Chicken embryo experimental design

Chicken embryos at 18 days of age were divided into four groups (Figure 3): NC group, Exo group, IBV group, and Exo+IBV group. The Exo and Exo+IBV groups received an injection of HD11_M1_-exo (50 μg per embryo) into the allantoic cavity, while the NC and IBV groups received an equal volume of PBS. The injection holes were sealed with wax. The embryos were returned to the incubator for an additional 24 hours. Subsequently, the IBV and Exo+IBV groups were inoculated with IBV strain (0.2 mL EID50 per embryo) into the allantoic membrane, while the NC and Exo groups received an equal volume of PBS. The embryos were then placed back into the chicken embryo incubator for further cultivation. At 24, 48, and 72 hours after inoculation, the embryos were dissected, and the clinical symptoms and pathological changes in each group were observed. The tracheas of the embryos were collected, and qPCR was used to detect changes in the levels of tracheal cell factors.

### Animal Experiment Design

During this experiment, the immunization method was via eye drops and nasal drops. The dosage of HD11_M1_-exo was 50 μg per chicken, the dosage of aluminum hydroxide adjuvant was 31.4 μg per chicken, the dosage of the 1313 adjuvant was 0.05 mL per chicken, and the dosage of the H120 IB attenuated vaccine was 0.2 mL EID50.

To evaluate the immunostimulatory potential of HD11_M1_-exo in vivo, day-old chicks (n=50) were randomly allocated into two experimental groups (n=25 per group) as illustrated in Figure 5. Birds received oculonasal administration of either HD11_M1_-exo (Exo group) or physiological saline (NC group) at days 1, 3, 5, and 14 post-hatch. Clinical status of birds was monitored daily throughout the experimental period. At 7, 14, 21, 28, and 35 days post-vaccination (**dpv**), five birds were randomly selected from each group at each time point for collection of lachrymal fluid, followed by euthanasia and subsequent tissue sampling. To evaluate the adjuvant effect of HD11_M1_-exo on immune responses to infectious bronchitis (IB) live attenuated vaccine, day-old chicks (n=100) were randomly allocated into four groups (n=25 per group) as detailed in Figure 8. H120 group: vaccinated with H120 strain alone; HD11+Exo group: vaccinated with H120 strain combined with HD11_M1_-exo; H120+AL group: vaccinated with H120 strain adjuvanted with aluminum hydroxide; and H120+1313 group: vaccinated with H120 strain adjuvanted with Montanide™ IMS 1313. Initial vaccination was performed at day 1 post-hatch via the oculonasal route. At 3 and 5 days post-vaccination (**dpv**), birds received a second and third dose of their respective adjuvants (HD11_M1_-exo, saline, aluminum hydroxide, or Montanide ™ IMS 1313) via the same route. A booster vaccination following the identical prime-vaccination protocol was administered at 14 dpv. Clinical status was monitored daily throughout the study period. At 7, 14, 21, 28, and 35 dpv, lachrymal fluid samples were collected from five randomly selected birds per group, after which the birds were euthanized for tissue collection.

To assess the protective efficacy of HD11_M1_-exo as an adjuvant during infectious bronchitis virus (IBV) challenge, day-old chicks were assigned to six experimental groups and NC group was housed separately under identical environmental conditions. Primary vaccination was administered at day 1 post-hatch (1 dpv) with the following regimens (Figure 12): the H120 group was inoculated only with the H120 vaccine; the HD11+exo group was inoculated with both the H120 vaccine and HD11_M1_-exo; the H120+AL group received the H120 vaccine with aluminum hydroxide adjuvant; the H120+1313 group received the H120 vaccine with the 1313 adjuvant; the NC group and IBV group were inoculated with an equal volume of physiological saline. At 14 dpv, all groups of chicks received a booster immunization following the same protocol as at 1 dpv. Seven days after the booster immunization, the IBV challenge was conducted; except for the NC group, each chick in the remaining groups was inoculated with 0.1 mL of EID50 IBV virus solution, while the NC group received 0.1 mL of PBS. Throughout the experiment, the clinical manifestations of the chicks in each group were observed daily, and on the fifth day post-infection (designated as 5 dpi), samples were collected.

### Real-time quantitative PCR

Quantitative reverse transcription PCR (RT-qPCR) analysis: Expression levels of LPS/TLR4 signaling pathway genes (CD14, TLR4, TRAM, TRAF3, IRF3, and CD80) were quantified in vitro using cultured cells. In the embryonated egg model, TNF-α mRNA expression was assessed in embryonic tracheal tissues at predetermined time points. In the in vivo study, mRNA expression of LPS/TLR4 pathway components was measured in tracheal tissues at 21 dpv. Additionally, expression of mucosal immune factor (TGF-β4) and co-stimulatory molecules (CD80 and CD86) was analyzed in tracheal and Harderian gland samples at various sampling points. Viral RNA load in tracheal tissues was determined at 5 dpi. Primer sequences used for gene amplification are listed in Table 1.

**Table 1.**
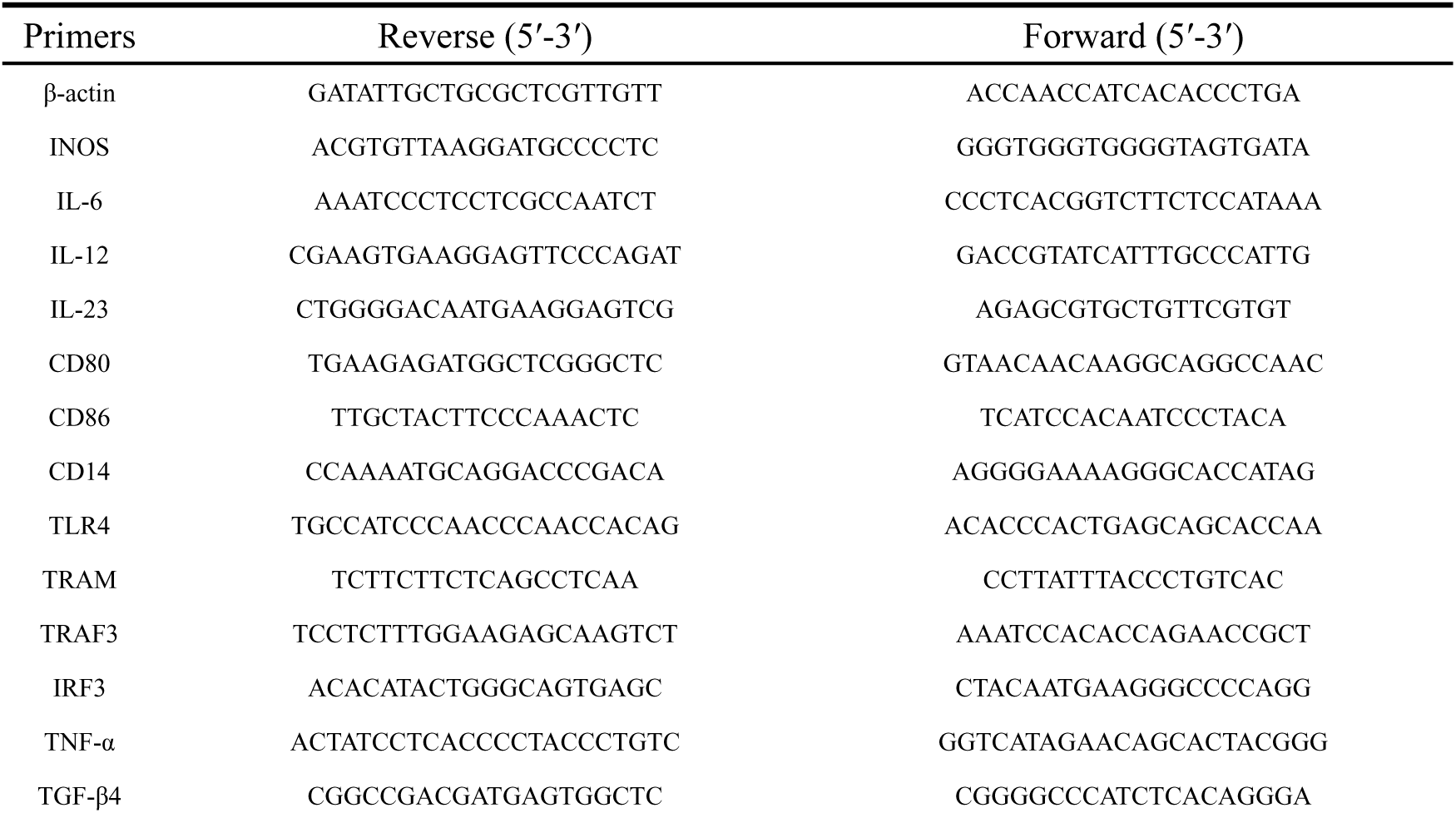
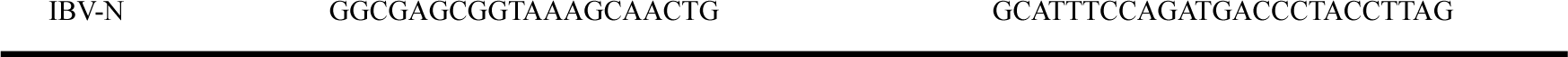
qPCR primer sequences.

### Tear fluid IgA antibody testing

Lachrymal fluid was collected from five birds per group at 7, 14, 21, 28, and 35 days post-primary vaccination (**dpv**). Briefly, lacrimation was induced by applying approximately 1 mg of molecular biology-grade sodium chloride crystals to the bilateral ocular surfaces. Tears were collected within 30-45 seconds post-stimulation using a calibrated micropipette. Samples were immediately transferred to sterile microcentrifuge tubes containing protease inhibitor cocktail (1:100 v/v) and stored at −20 °C until analysis. Tear IgA concentrations were determined using a commercial chicken-specific IgA ELISA kit according to the manufacturer’s instructions.

### Serum antibody level testing

At 7, 14, 21, 28, and 35 days post-prime vaccination (**dpv**), blood samples were collected from birds in each experimental group via wing vein puncture. Samples were allowed to clot at room temperature for 2 h and subsequently centrifuged at 5000 × g for 15 minutes to obtain serum fractions. Serum antibody titers against avian infectious bronchitis virus (IBV) were determined using a commercially available indirect ELISA kit (Company, Location) per manufacturer’s specifications.

### Histopathological observations

Collect tracheal tissues from different groups of chickens in the immunological detoxification protection experiment for histopathological analysis. The tissues are fixed in 4% formalin solution. After dehydration with ethanol, the paraffin-embedded tracheal tissues are sectioned to a thickness of 5 microns and stained with hematoxylin and eosin (H&E). Following staining, the sections are dehydrated with absolute ethanol and washed with xylene for transparency. Finally, transparent sections are mounted with neutral resin and examined under a microscope to evaluate the lesions in each group. The observations are photographed and recorded.

### Statistical Analysis

The experimental data are presented as the means ± standard deviation. Statistical analyses were performed using GraphPad Prism (9.5.1), with comparisons made using a one - way analysis of variance or multiple comparisons. Significance levels are indicated as follows: Different letters indicate that the differences are significant at the *p* < 0.05 level. Specifically, treatment A (a) does not differ significantly from treatment B (ab), but it does differ significantly from treatment C (b); treatment B (ab) also does not differ significantly from treatment C (b). Data visualization was accomplished using GraphPad Prism (9.5.1) software.

## Results

### Isolation and characterization of exosomes derived from M1-type macrophages

To determine whether macrophages polarize to the M1 type after stimulation with LPS, qPCR was used to detect typical markers of M1 type in macrophages from the stimulated group (LPS group) and the control group (NC group) after 24 hours of stimulation. The results indicated that, compared to the NC group, the expression levels of INOS, IL-6, IL-12, IL-23, CD80, and CD86 in the LPS group were significantly elevated (*P* < 0.05) (Figure 1, A), demonstrating that macrophages had indeed polarized to the M1 type following LPS stimulation (***HD11_M1_-exo***).

**Figure 1.**
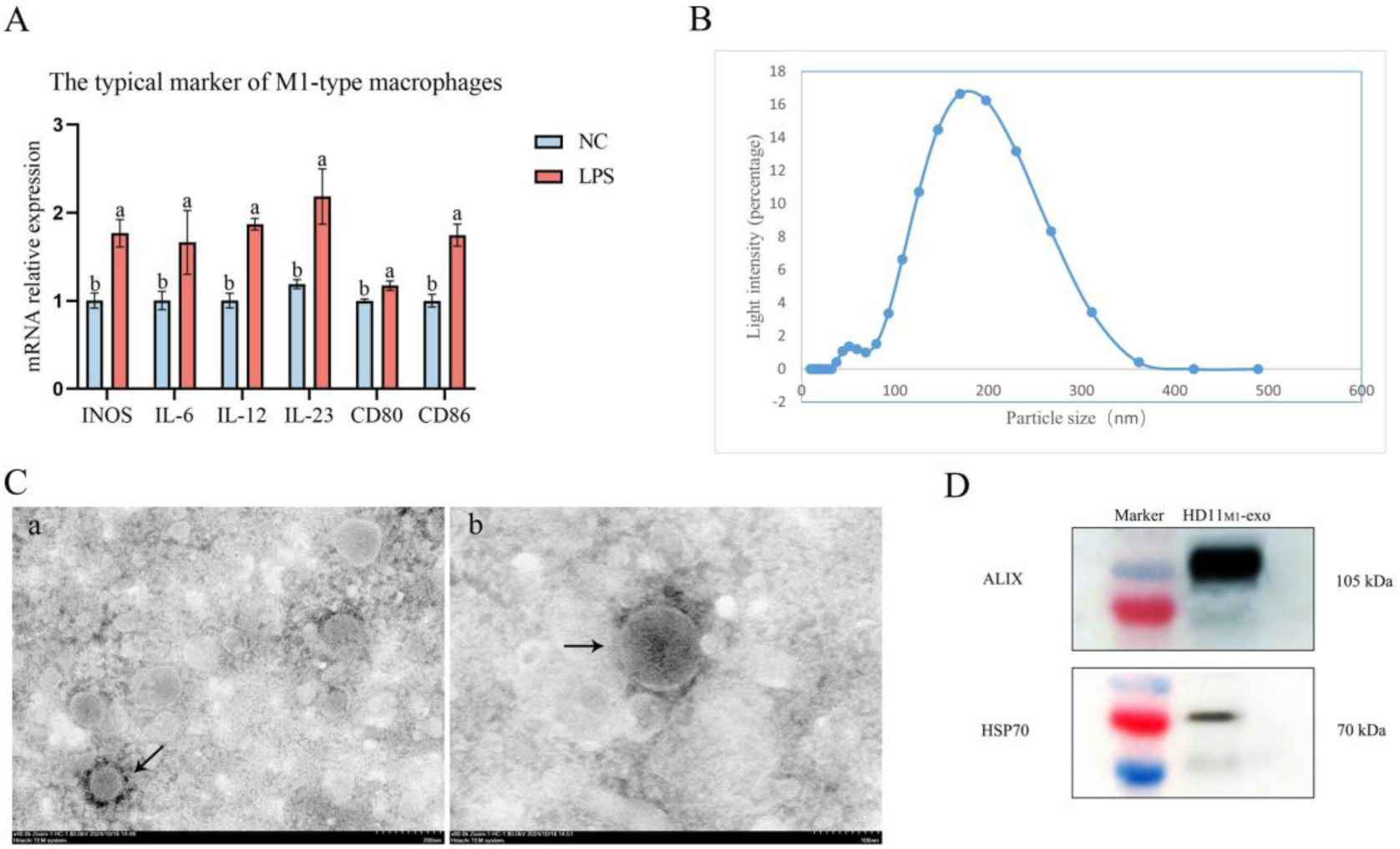
Characterization of HD11_M1_-exo. (A) Quantification of mRNA levels of M1 macrophage-specific markers. (B) Nanoparticle tracking analysis of HD11_M1_-exo. (C) Transmission electron microscopy (TEM) images of HD11_M1_-exo (black arrows) at magnifications of (a) 40,000× and (b) 80,000×. (D) Western blot detection of exosomal marker proteins ALIX and HSP70. Different superscript letters denote significant differences (*p* < 0.05).

The HD11_M1_-exo were separated using ultrahigh-speed centrifugation, and their characterization was identified through transmission electron microscopy, particle size analysis, and Western Blot. The experimental results indicate that HD11_M1_-exo have an intact structure and exhibit cup-shaped or spherical forms, with particle sizes primarily distributed between 30 and 200 nm, in accordance with the characteristics of exosomes (Figure 1, B C). Western Blot analysis detected positive signals for the typical exosomal markers HSP70 (70 kDa) and ALIX (105 kDa) in HD11_M1_-exo (Figure 1, D). These results suggest that successful separation of HD11_M1_-exo has been achieved.

### HD11_M1_-exo enhance the immune activity of macrophages through activate the LPS/TLR4 signaling pathway

After 24 hours of stimulation with HD11_M1_-exo, compared to the NC group, macrophages in the exosome group showed a significant increase in their phagocytic capacity (*P* < 0.05) (Figure 2 A). Additionally, after 24 hours of HD11_M1_-exosome stimulation, the expression levels of key genes in the LPS/TLR4 signaling pathway, including CD14, TLR4, TRAM, TRAF3, IRF3, and CD80, were significantly upregulated in the exosome group (*P* < 0.05) (Figure 2 B), with the mechanism of action illustrated in Figure 2 C. These experimental results indicate that HD11_M1_-exo can activate the LPS/TLR4 signaling pathway in macrophages, enhancing their phagocytic ability and thus boosting their immune activity.

**Figure 2.**
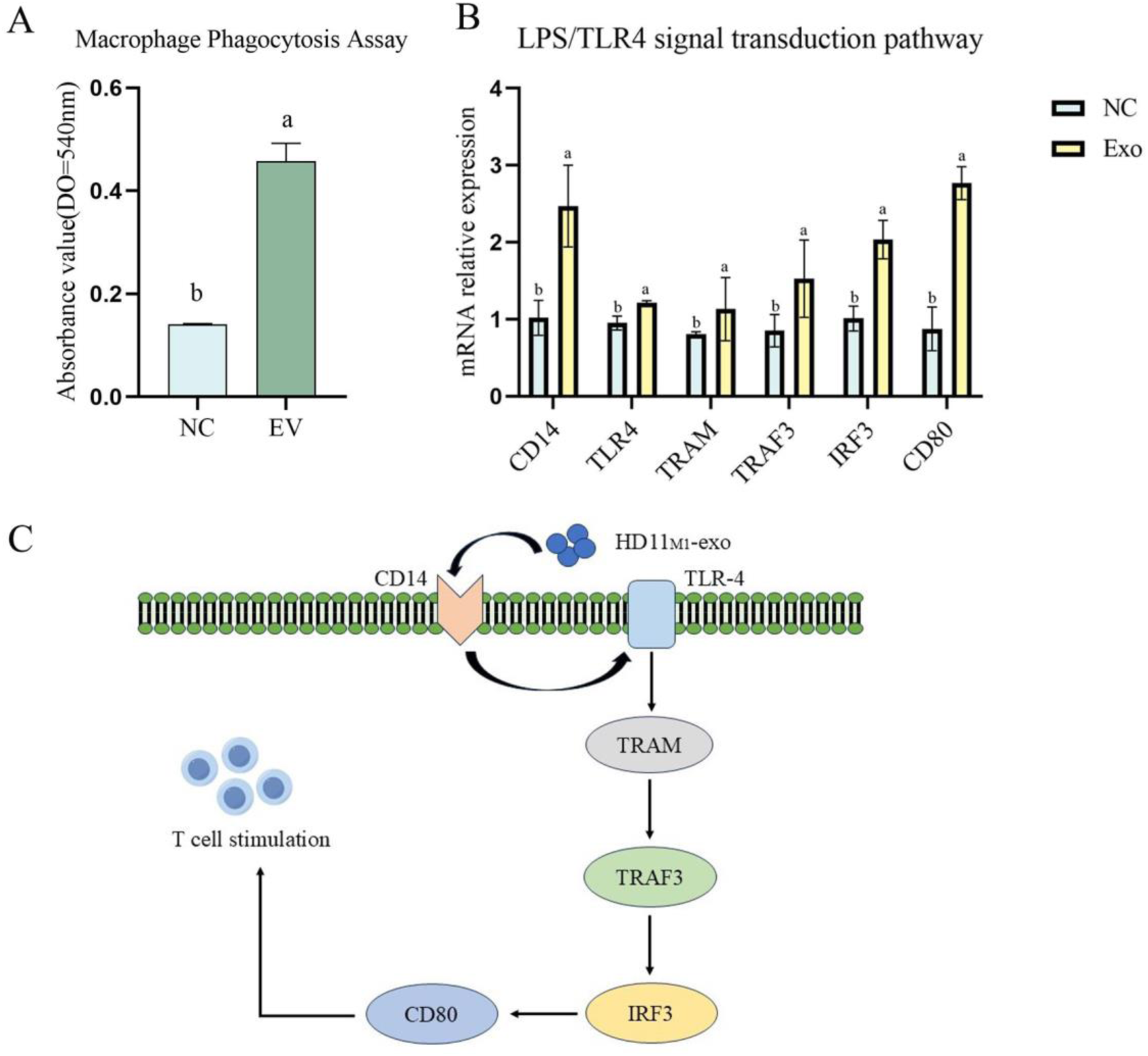
Effects of HD11_M1_-derived exosomes on macrophage function after 24 h stimulation. (A) Phagocytic capacity of macrophages assessed by neutral red uptake assay. (B) Relative mRNA expression of key genes in the LPS/TLR4 signaling pathway. (C) Schematic illustration of HD11_M1_-derived exosome-mediated macrophage activation through the LPS/TLR4 signaling pathway. Different superscript letters denote significant differences (*p* < 0.05).

### HD11_M1_-exo pretreatment enhanced antiviral resistance to IBV infection in embryonated chicken eggs

The experimental design of HD11_M1_-exo resisting IBV infection in chicken embryos is shown in Figure 3.

**Figure 3.**
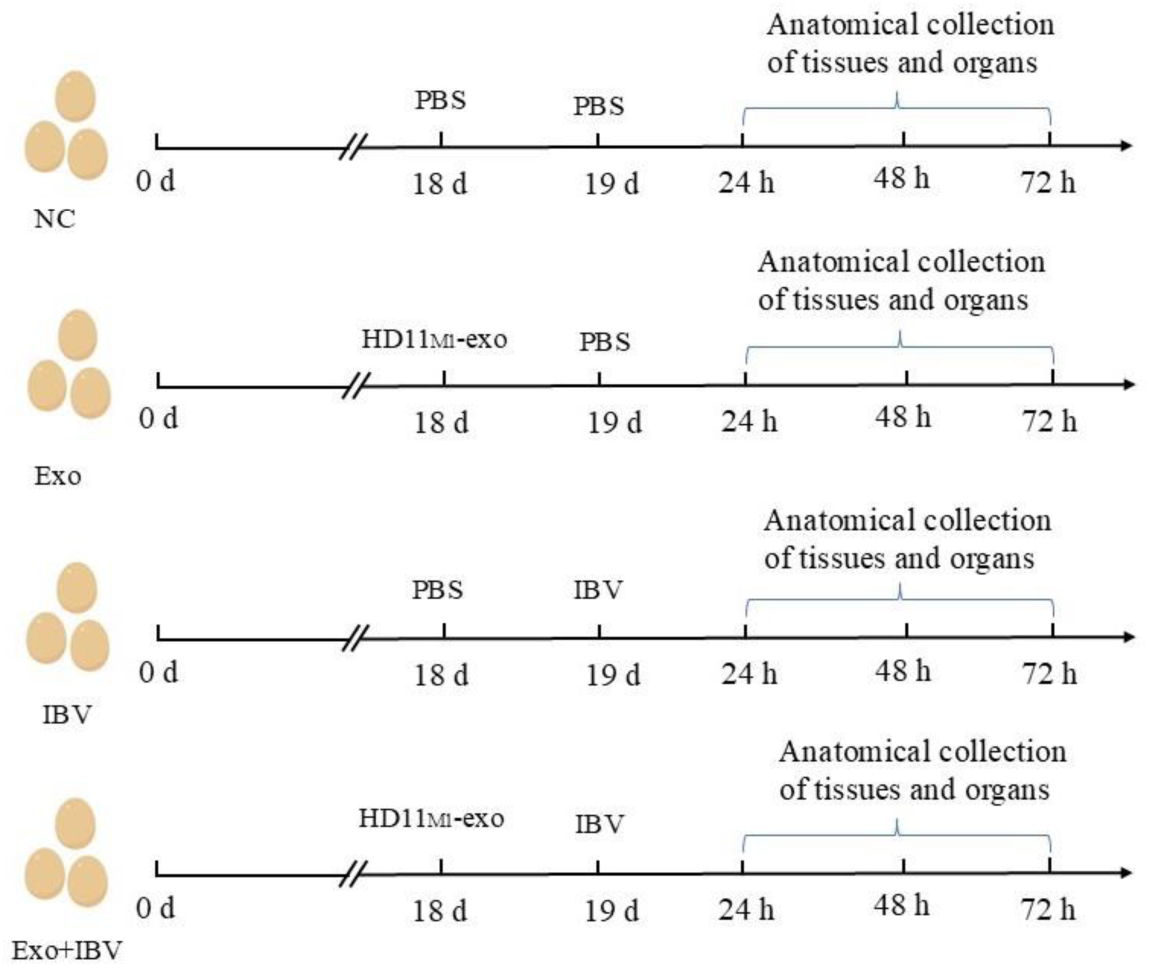
Chick embryo experiment design.

After stimulating the chicken embryos with HD11_M1_-exo for 24 hours, IBV was introduced. At 24 hours, 48 hours, and 72 hours post-infection, the IBV group showed 2, 1, and 1 chicken embryos, respectively, exhibiting symptoms of IBV infection (marked by blue arrows), while the NC group, Exo group, and Exo+IBV group showed no signs of lesions (Figure 4 A). Using qPCR to assess TNF-α expression in the tracheas of chicken embryos at 24, 48, and 72 hours post-infection revealed that the expression level of the TNF-α gene in the tracheal tissue of the Exo group was significantly increased compared to the NC group (P<0.05). Furthermore, compared to the IBV group, the TNF-α gene expression level in the tracheal tissue of the Exo+IBV group was also significantly elevated (P<0.05) (Figure 4 B). The results indicate that HD11_M1_-exo enhances the natural immune response of chicken embryos, improving their resistance to IBV.

**Figure 4.**
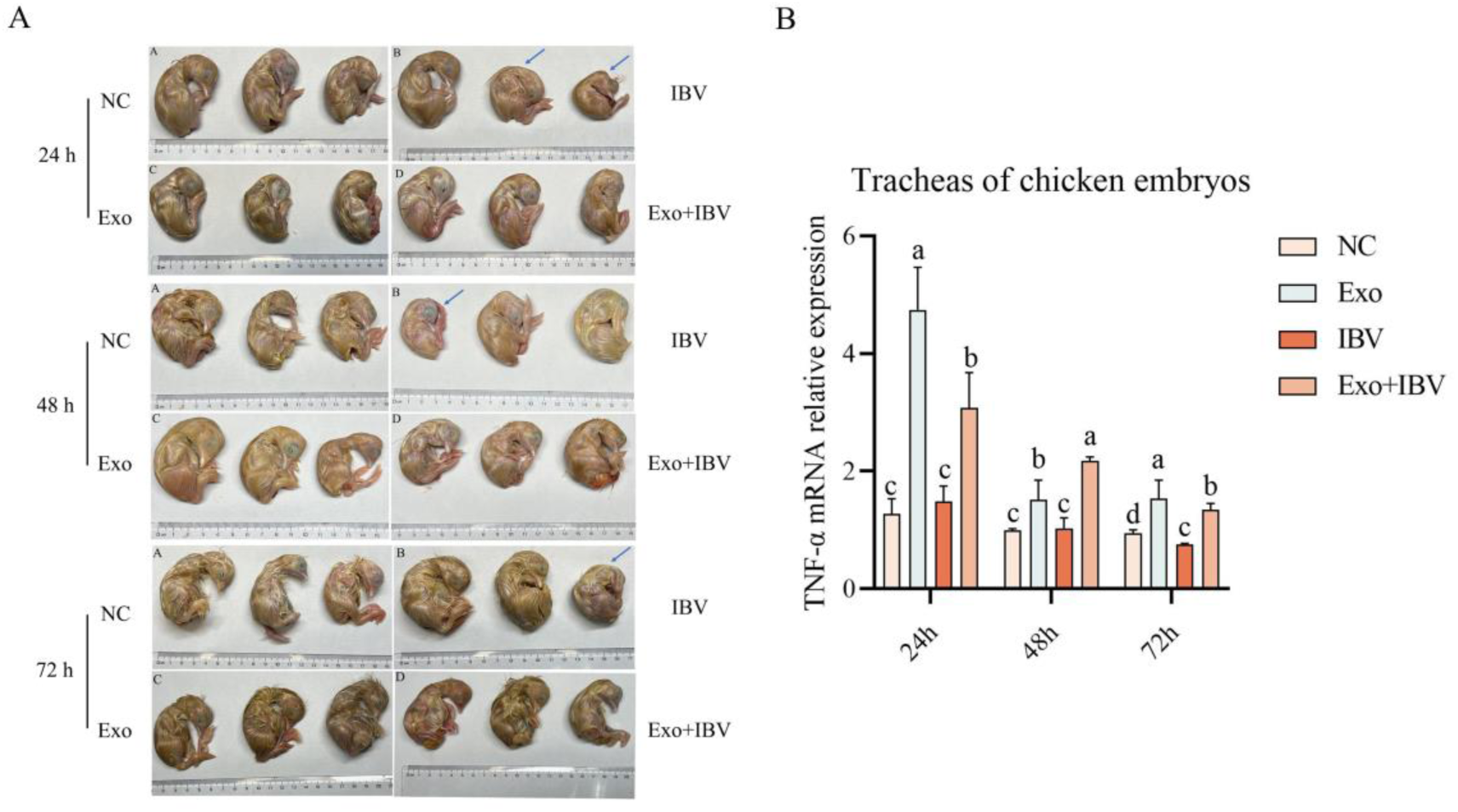
Embryonated egg challenge model. Embryos were pretreated with HD11_M1_-derived exosomes for 24 h before IBV challenge. (A) Gross pathology of IBV-infected embryos at 24, 48, and 72 h post-infection (hpi). Arrows indicate characteristic IBV-induced lesions. (B) Relative TNF-α mRNA expression in tracheal tissues. Different superscript letters denote significant differences (*p* < 0.05).

### HD11_M1_-exo potentiate cellular immune responses in chickens

The experimental design for the immune activation induced by HD11_M1_-exo in chickens is shown in Figure 5.

**Figure 5.**
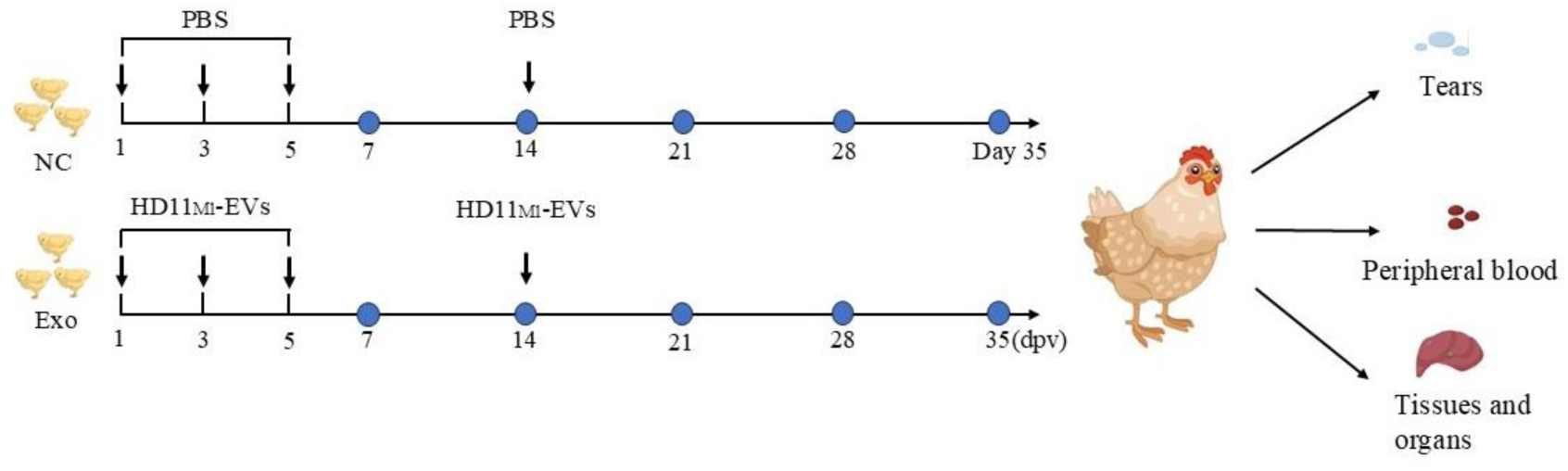
Immunoactivation experimental design in animals. Note: Blue dots indicate dissection time points.

Quantitative PCR analysis was performed to assess the mRNA expression of co-stimulatory molecules CD80 and CD86 in tracheal and Harderian gland tissues from HD11_M1_-Exos-treated chickens at 7, 14, 21, 28, and 35 days post-vaccination (**dpv**). Expression of both CD80 and CD86 was significantly upregulated (*P* < 0.05) in both tissues from the HD11_M1_-Exos group compared to the negative control group at all examined time points (Figures 6A-D). This consistent upregulation throughout the 35-day observation period suggests sustained immunomodulatory effects of HD11_M1_-Exos on co-stimulatory molecule expression.

**Figure 6.**
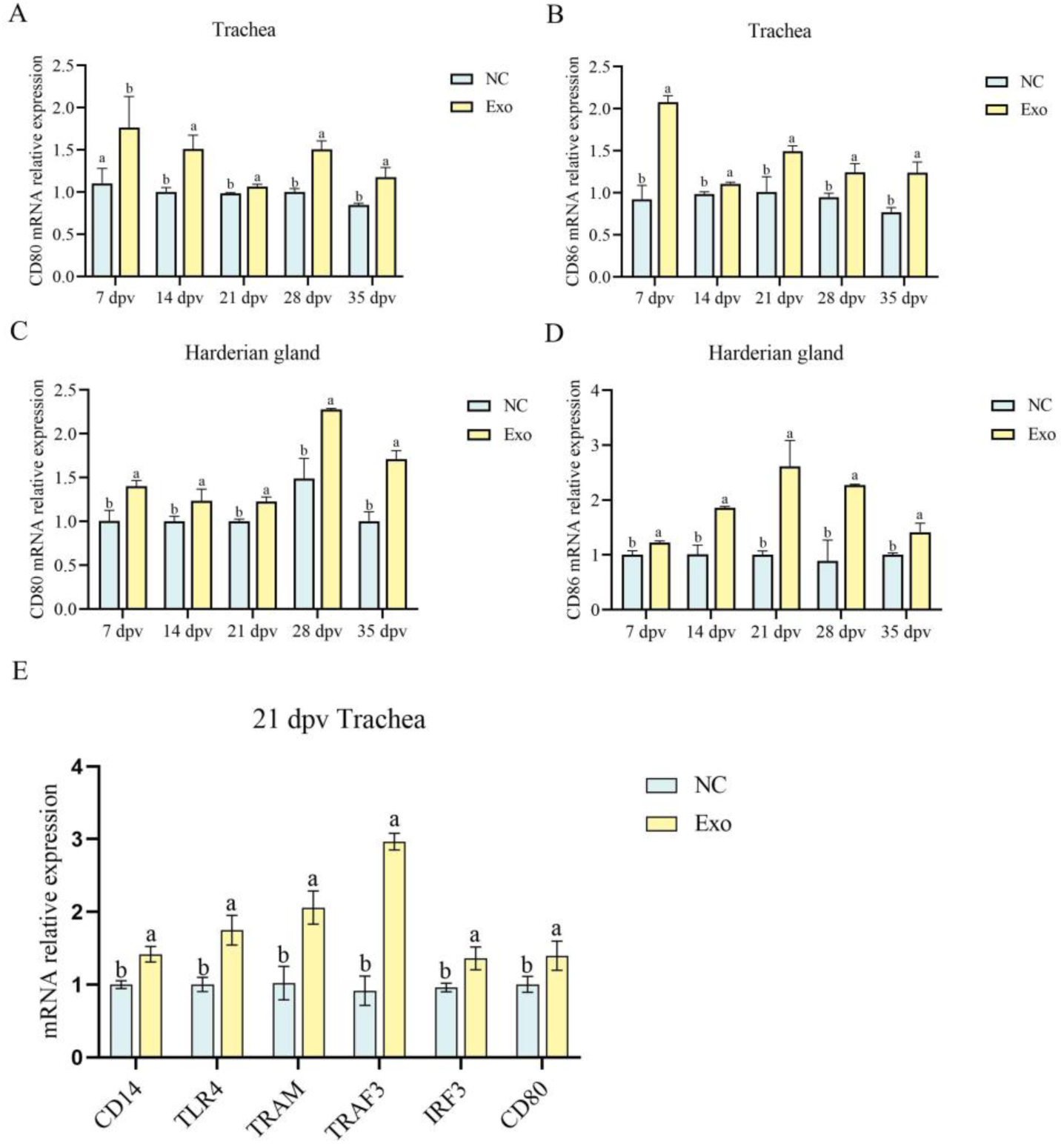
A. mRNA levels of the CD80 gene in the trachea. B. mRNA levels of the CD86 gene in the trachea. C. mRNA levels of the CD80 gene in the Harderian gland. D. mRNA levels of the CD86 gene in the Harderian gland. E. mRNA levels of key genes in the LPS/TLR4 signaling pathway in the trachea at 21 dpv. Different superscript letters denote significant differences (*p* < 0.05).

To elucidate the molecular mechanisms underlying the immunostimulatory effects of HD11_M1_-exo in chickens, the expression of genes involved in the LPS/TLR4 signaling pathway was evaluated in tracheal tissues from immunized birds using quantitative PCR. Analysis revealed significant upregulation (*P* < 0.05) of key components of the LPS/TLR4 signaling cascade, including CD14, TLR4, TRAM, TRAF3, IRF3, and CD80, in the Exo group compared to the NC group (Figure 6E).

These findings demonstrate that HD11_M1_-exo treatment significantly upregulates expression of co-stimulatory molecules CD80 and CD86 in tracheal and Harderian gland tissues, activates the LPS/TLR4 signaling pathway, and potentiates cellular immune responses in chickens.

### HD11_M1_-exo augments mucosal immunity in the respiratory tract of chickens

Using qPCR and a chicken immunoglobulin A (IgA) kit (ELISA), we detected the levels of the mucosal immune-related gene TGF-β4 in the trachea and IgA in the tears of experimental chickens at 7 dpv, 14 dpv, 21 dpv, 28 dpv, and 35 dpv. The experimental results showed that, compared to the NC group, the transcription level of the mucosal immune-related gene TGF-β4 in the trachea of the Exo group was significantly elevated at each time point (*P*<0.05) (Figure 7 A). Additionally, the level of chicken immunoglobulin IgA in the tears of the Exo group was significantly elevated at each time point compared to the NC group (*P*<0.05) (Figure 7 B). These results suggest that stimulation with HD11_M1_-exo can enhance the transcription level of the mucosal immune-related gene TGF-β4 and the level of chicken immunoglobulin IgA, thereby improving the mucosal immunity in chickens.

**Figure 7.**
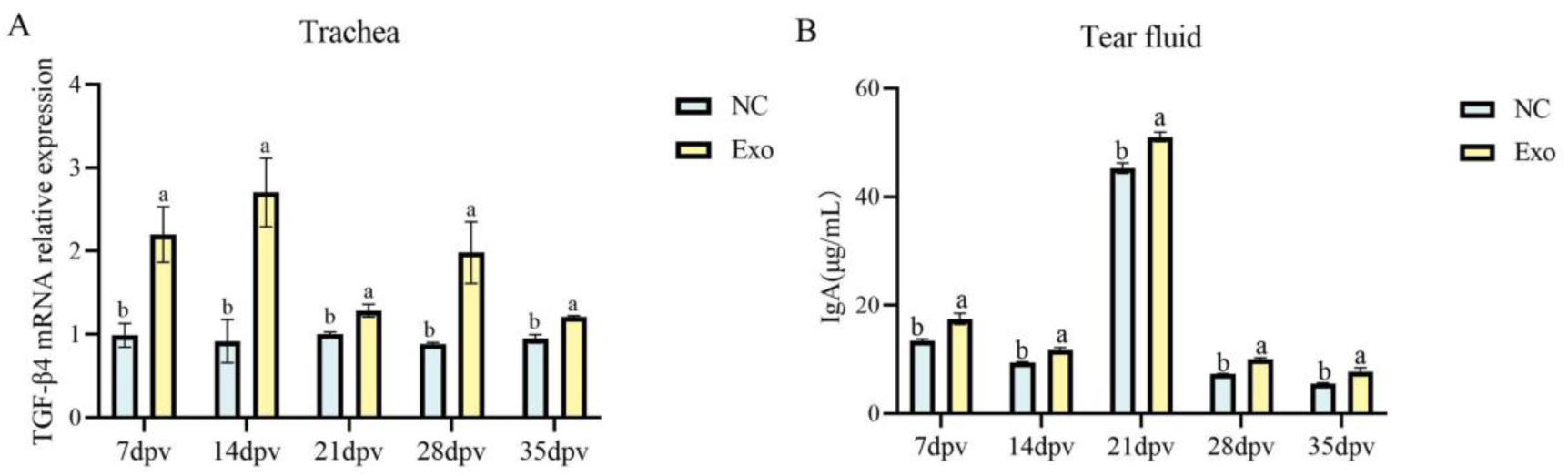
Mucosal immune level detection. A. mRNA levels of the mucosal immune-related gene TGF-β4 in the trachea. B. Levels of chicken immunoglobulin IgA in tears. Different superscript letters denote significant differences (*p* < 0.05).

### HD11_M1_-exo enhance vaccine H120-induced cellular immune responses in chickens

The experimental design for immunological activation induced by the combined use of IB vaccines in chickens is shown in Figure 8.

**Figure 8.**
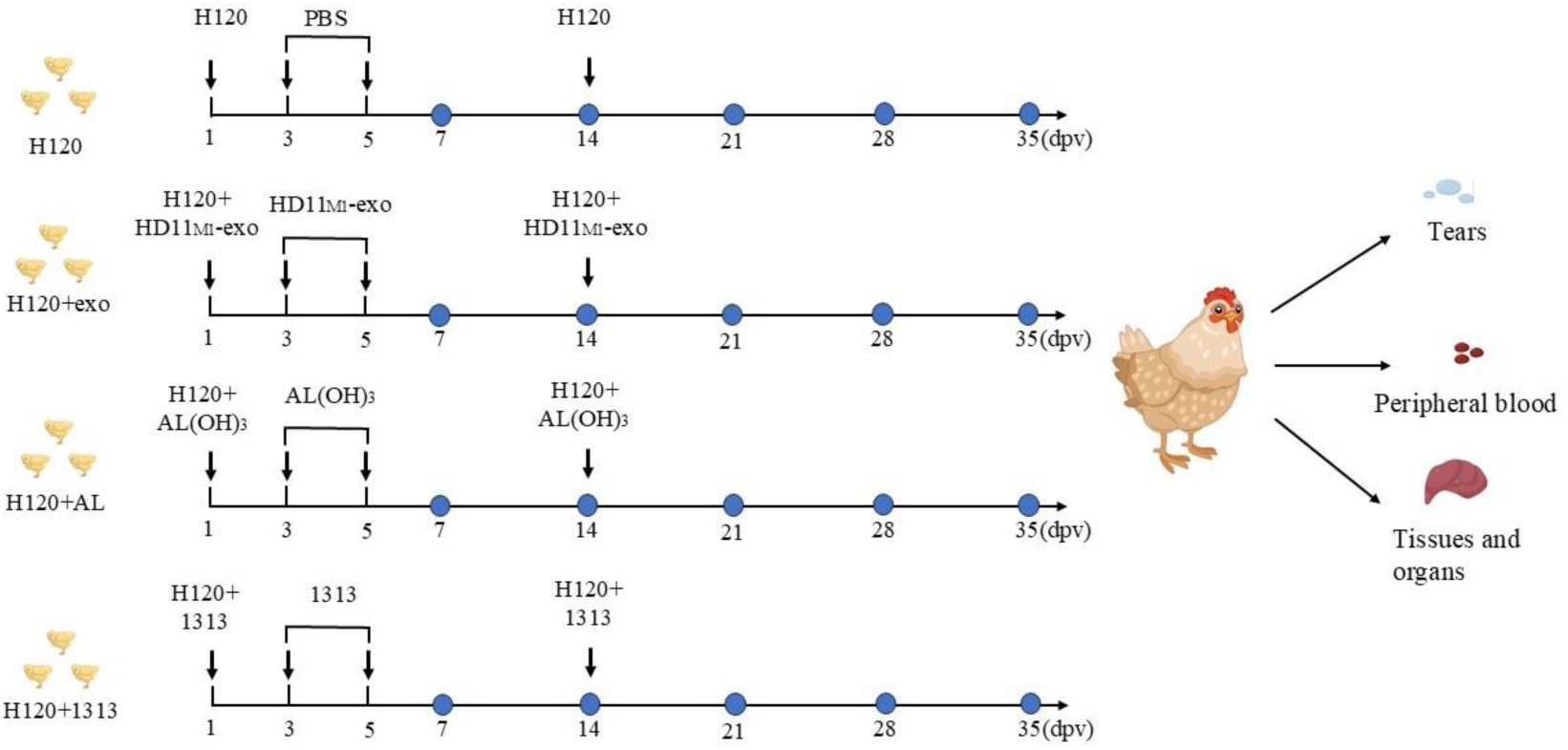
Animal experiment design. Note: Blue circles indicate dissection time points.

In this experiment, qPCR was used to detect the transcription levels of the co-stimulatory factors CD80 and CD86 in the tracheas and Harderian glands of each group of experimental chickens at 7 dpv, 14 dpv, 21 dpv, 28 dpv, and 35 dpv. The experimental results indicated that, compared to the use of the vaccine alone, the combination of HD11_M1_-exo with the vaccine significantly increased the transcription levels of CD80 and CD86 in the tracheas and Harderian glands of immunized chickens. Furthermore, the effects were more pronounced at 7 dpv and 28 dpv compared to the H120+AL group and the H120+1313 group (*P*<0.05) (Figure 9 A-D). In summary, the combination of HD11_M1_-exo with the IB attenuated vaccine H120 significantly enhances the transcription levels of the co-stimulatory factors CD80 and CD86 in the tracheas and Harderian glands of immunized chickens, indicating that HD11_M1_-exo can improve the cellular immune response induced by the vaccine.

**Figure 9.**
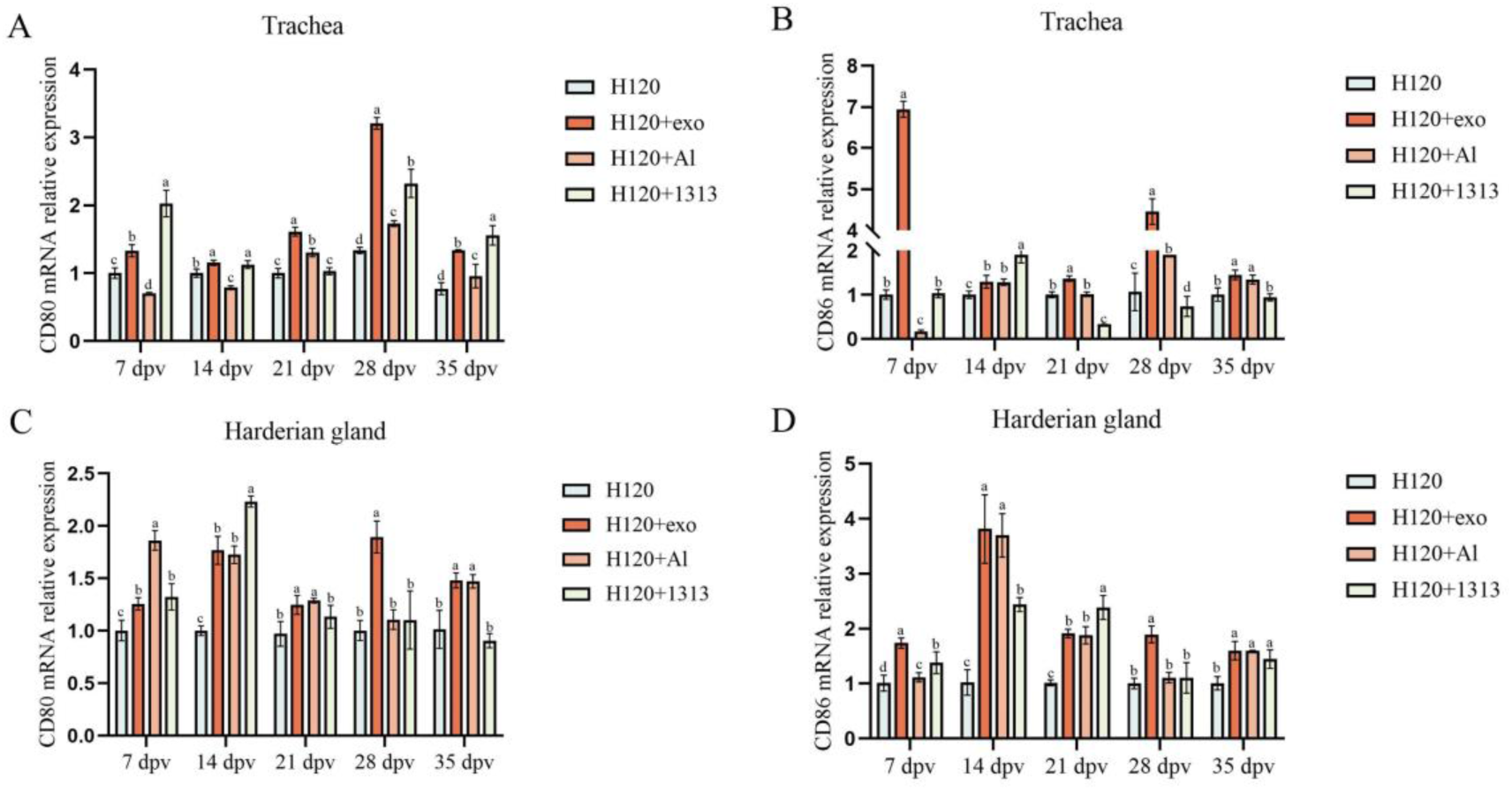
Immune co-stimulatory marker expression in chickens. (A) Relative mRNA expression of the CD80 gene in tracheal tissues. (B) Relative mRNA expression of the CD86 gene in tracheal tissues. (C) Relative mRNA expression of the CD80 gene in Harderian gland tissues. (D) Relative mRNA expression of the CD86 gene in Harderian gland tissues. Different superscript letters denote significant differences (*p* < 0.05).

Expression of co-stimulatory molecules CD80 and CD86 in tracheal and Harderian gland tissues was assessed by quantitative PCR at 7, 14, 21, 28, and 35 days post-vaccination (**dpv**) across all experimental groups. The HD11_M1_-exo adjuvanted vaccine significantly upregulated CD80 and CD86 mRNA levels in both tissues compared to vaccination with H120 alone. This enhancement was particularly evident at 7 and 28 dpv, where the H120+exo group significantly higher expression than both the H120+AL and H120+1313 groups (*P* < 0.05) (Figures 9A-D). Collectively, these data demonstrate that HD11_M1_-exo, when combined with the attenuated infectious bronchitis vaccine H120, potentiates the expression of key co-stimulatory molecules in mucosal-associated lymphoid tissues, suggesting enhanced vaccine-induced cellular immunity compared to conventional adjuvant formulations.

### HD11_M1_-exo significantly elevates vaccine H120-induced humoral immunity in chickens

Serum IgY antibody levels against infectious bronchitis virus were quantified using an indirect enzyme-linked immunosorbent assay (ELISA) at 7, 14, 21, 28, and 35 days post-vaccination (dpv). Administration of HD11_M1_-exo in combination with the attenuated infectious bronchitis vaccine H120 significantly enhanced vaccine-induced antibody responses throughout the observation period. This enhancement was particularly pronounced at 14 dpv, when the HD11+exo group exhibited significantly higher antibody titers compared to both the H120+AL and H120+1313 groups (*P* < 0.05) (Figure 10). These findings demonstrate that HD11_M1_-exo effectively potentiates the humoral immune response elicited by the H120 vaccine.

**Figure 10.**
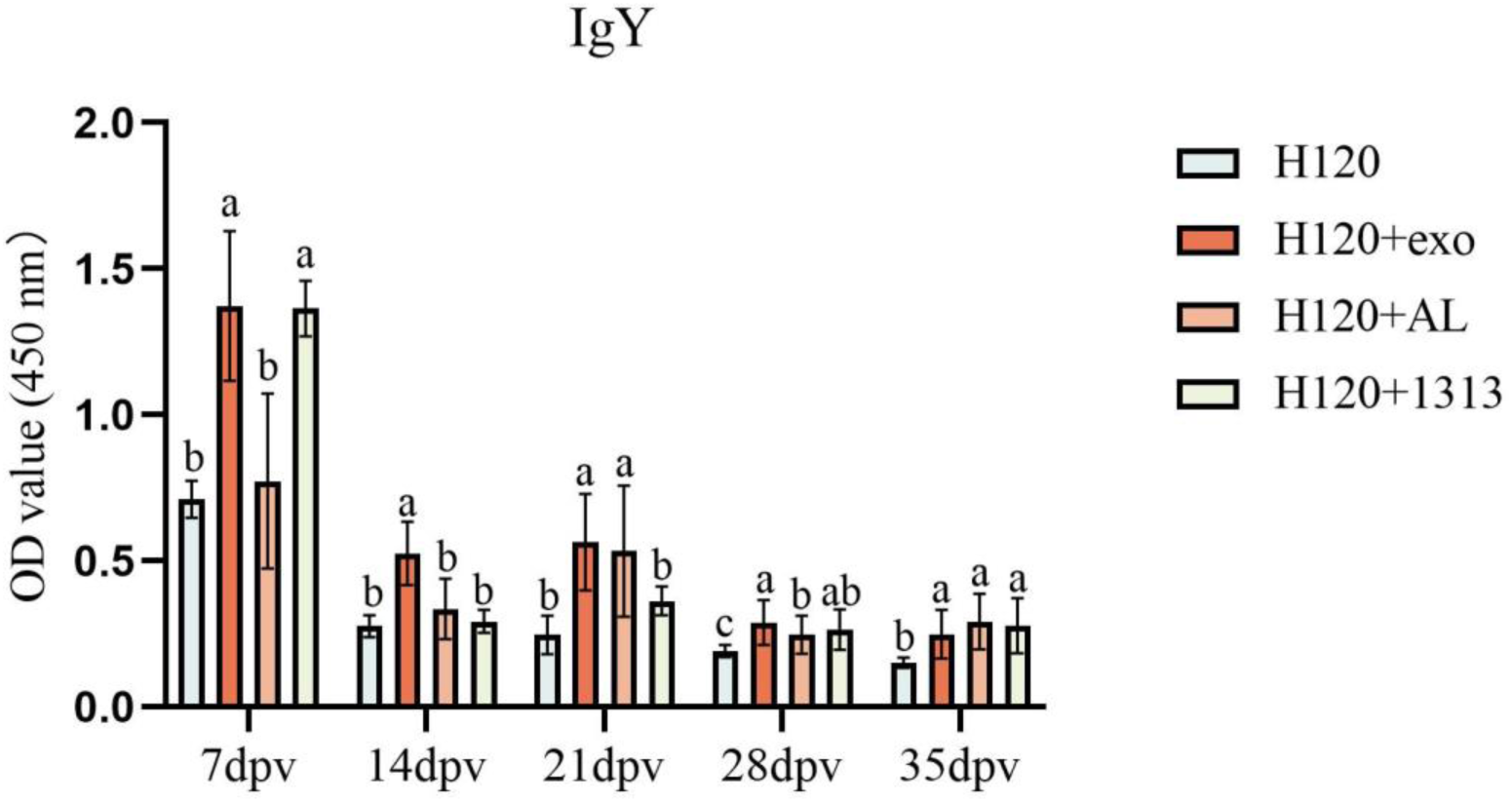
Levels of IgY antibodies in the serum of experimental chickens’ peripheral blood. Different superscript letters denote significant differences (*p* < 0.05).

### HD11_M1_-exo enhance vaccine H120-induced mucosal IgA production in the avian respiratory tract

HD11_M1_-exo was administered in combination with the attenuated infectious bronchitis vaccine H120. Lachrymal fluid and tracheal tissue samples were collected at 7, 14, 21, 28, and 35 days post-vaccination (**dpv**). Secretory IgA concentrations in lachrymal fluid were quantified using enzyme-linked immunosorbent assay (ELISA), while TGF-β4 mRNA expression in tracheal tissues was measured by quantitative PCR. The H120+exo groups formulation significantly elevated both lachrymal IgA concentrations and tracheal TGF-β4 expression compared to other groups (*P* < 0.05) (Figures 11A, B). These results demonstrate that HD11_M1_-exo markedly enhances vaccine-induced mucosal immune responses at respiratory surfaces.

**Fig 11.**
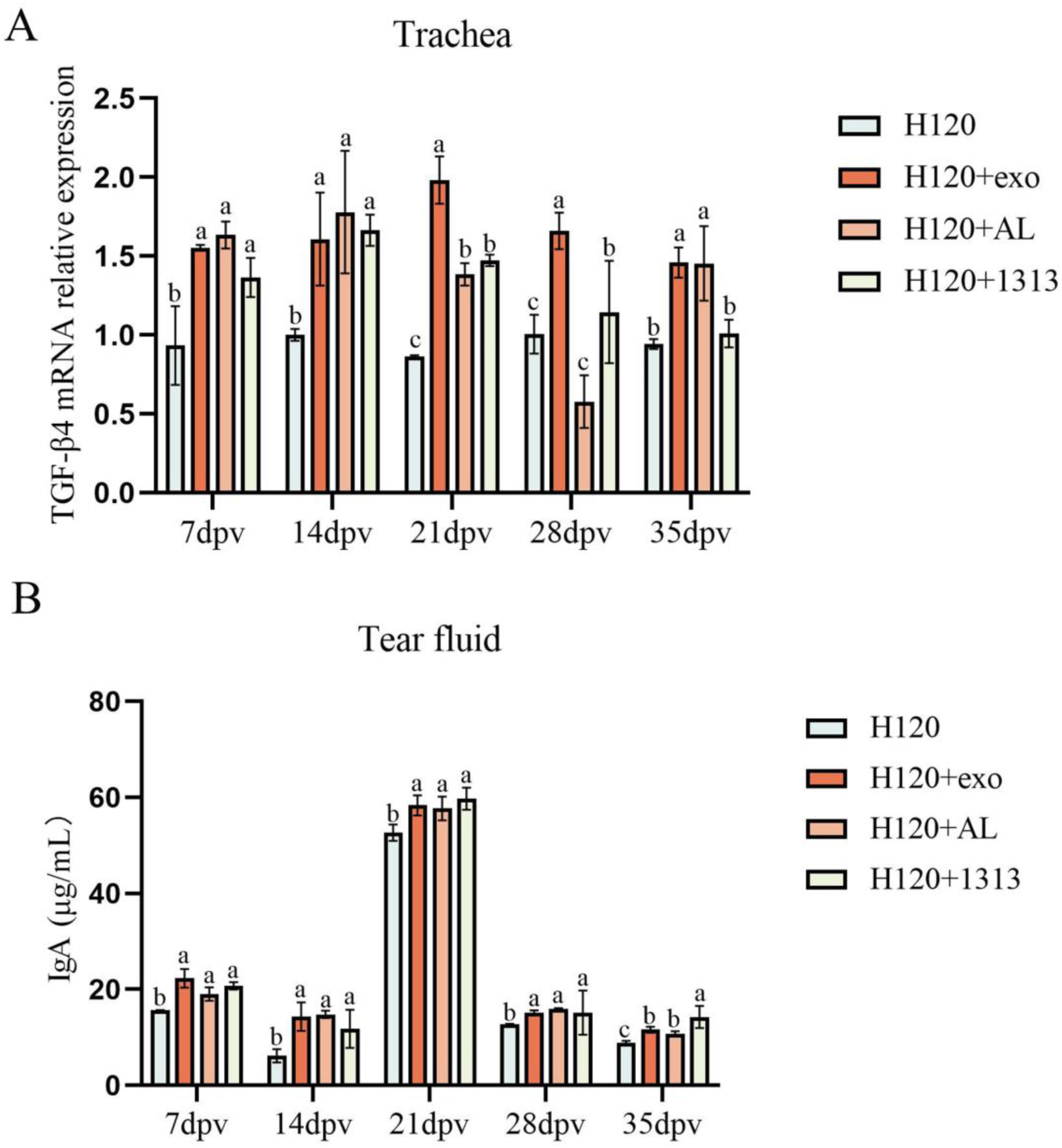
Mucosal immunity assessment. (A) Relative mRNA expression of mucosal immunity-associated TGF-β4 in tracheal tissues. (B) IgA concentrations in tear samples. Different superscript letters denote significant differences (*p* < 0.05).

### HD11_M1_-exo Enhance Protective Efficacy of H120 Vaccine against IBV Challenge

The experimental design for the combined use of HD11_M1_-exo and IB vaccine to resist IBV infection in chickens is shown in Figure 12.

**Figure 12.**
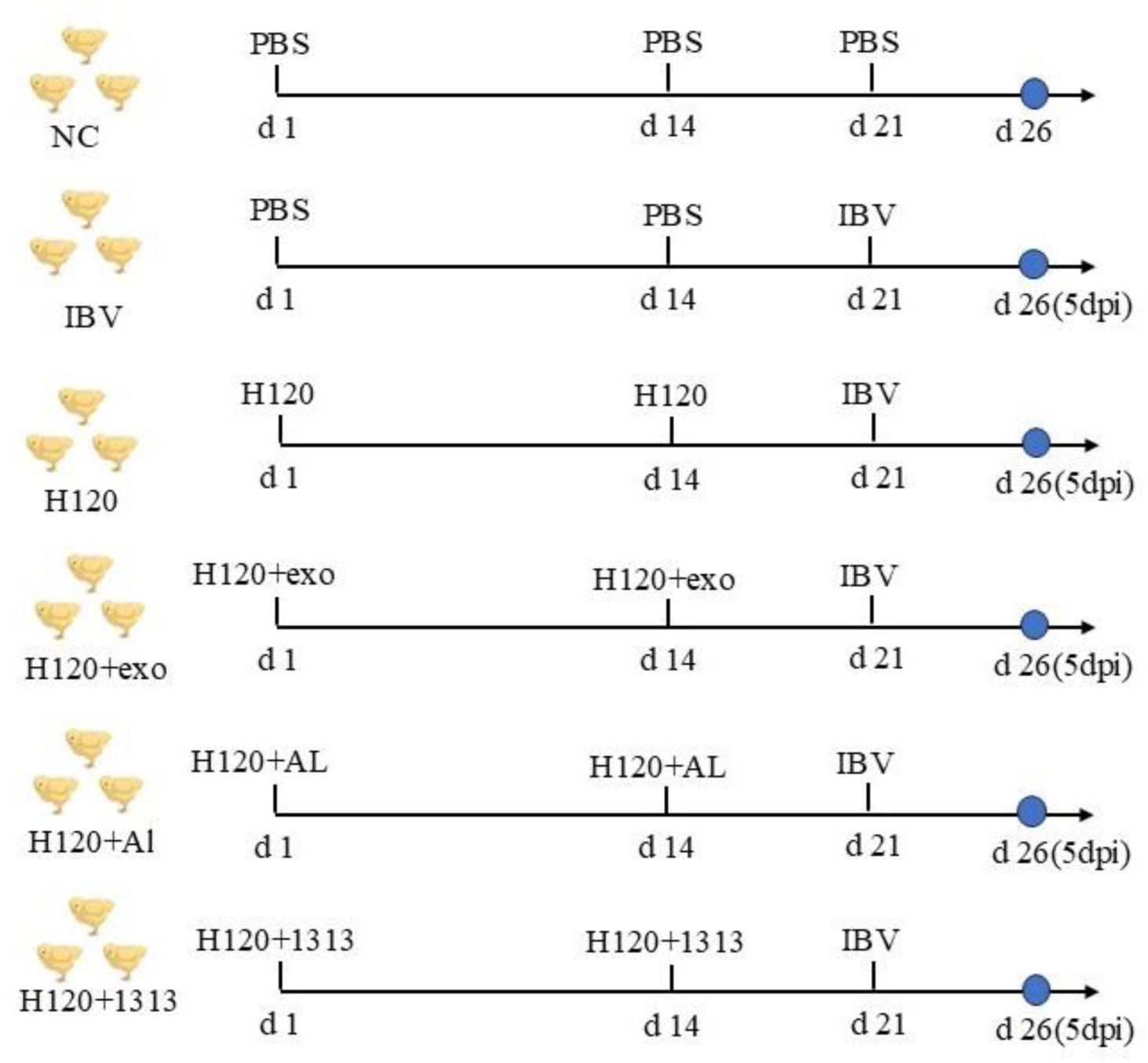
Design of animal immune response protection experiments against toxins. Note: Blue dots indicate dissection time points.

Viral load in tracheal tissues at 5 days post-infection (**dpi**) was assessed by quantitative PCR measuring relative expression of the IBV nucleocapsid (N) gene. Analysis revealed significantly reduced viral burdens in tracheal tissues from chickens in both the HD11+exo and H120+Al groups compared to the H120 group (*P* < 0.05) (Figure 13A). In contrast, viral loads in the H120+1313 group did not differ significantly from those in the H120 group. These findings demonstrate that HD11_M1_-exo adjuvantation of the H120 vaccine effectively reduces viral replication in the respiratory tract following challenge with infectious bronchitis virus.

**Figure 13.**
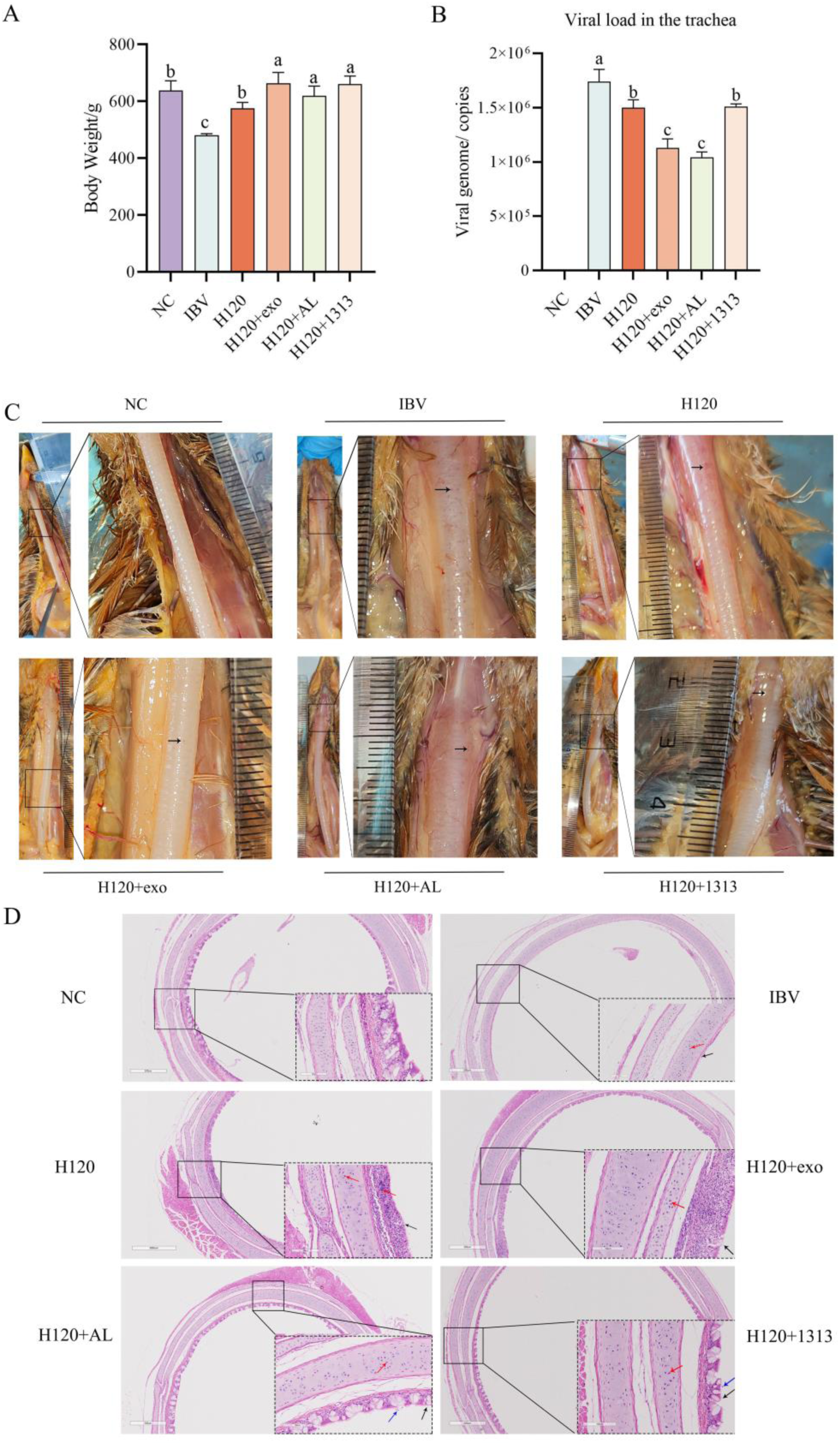
Immunotoxicity assessment in chickens at 5 days post-infection (**dpi**). (A) Body weight measurements across experimental groups. (B) Viral load in tracheal tissues. (C) Gross pathological examination of trachea; black arrows denote hemorrhagic foci. (D) Histopathological features of tracheal tissues; black arrows: ciliary loss; red arrows: lymphocyte infiltration; blue arrows: epithelial cell shedding. Different superscript letters denote significant differences (*p* < 0.05).

At 5 dpi, the body weight of each group of chickens was measured and subjected to statistical analysis. The results showed that the body weights of the NC group, H120 group, HD11+exo group, H120+AL group, and H120+1313 group were significantly higher than that of the IBV group (*P*<0.05) (Figure 13 B). Additionally, the body weights of the HD11+exo group, H120+AL group, and H120+1313 group were all significantly higher than that of the H120 group (*P*<0.05). Body weights were recorded and analyzed at 5 days post-infection (**dpi**). All vaccinated groups (H120, HD11+exo, H120+AL, and H120+1313) and the NC group exhibited significantly higher body weights compared to the unvaccinated, challenged IBV group (*P* < 0.05) (Figure 13B). Moreover, chickens in the adjuvanted vaccine groups (HD11+exo, H120+AL, and H120+1313) maintained significantly higher body weights than those receiving the H120 vaccine alone (*P* < 0.05). Necropsy examinations performed at 5 dpi revealed normal tracheal mucosa without lesions in the NC group, whereas numerous petechial hemorrhages were observed in tracheal tissues from the IBV group. The H120 group displayed moderate hemorrhagic lesions, while all adjuvanted groups (HD11+exo, H120+AL, and H120+1313) showed reduced tracheal pathology (Figure 13C). These findings confirm successful establishment of the challenge model and demonstrate that HD11_M1_-exo, when used as an adjuvant with the H120 vaccine, mitigates IBV-induced pathology and weight loss in infected chickens.

The pathological examination of the tracheal tissue samples indicates that the NC group of experimental chickens exhibited intact histological structure with no pathological changes. The IBV group displayed complete loss of cilia (as indicated by the black arrows), with the cell nuclei exhibiting deep staining and significant infiltration of lymphocytes (as shown by the red arrows). The H120 vaccine group showed a more pronounced loss of cilia (as indicated by the black arrows), with similarly deep-stained nuclei, but a comparatively lower number of lymphocyte infiltrates (as shown by the red arrows). The HD11+exo group showed only a small amount of cilia loss (as indicated by the black arrows), deep-stained nuclei (as shown by the red arrows), but no significant lymphocyte infiltration was observed. Both the H120+AL group and the H120+1313 group demonstrated slight cilia loss (as indicated by the black arrows), deep-stained nuclei (as shown by the red arrows), accompanied by some sloughing of epithelial cells (as indicated by the blue arrows) (Figure 13 D). In summary, compared to the H120 group, the combination of HD11_M1_-exo with the H120 vaccine can mitigate the pathological damage caused by IBV infection in the tracheas of chickens.

## Discussion

Immune cell-derived exosomes are currently the most extensively studied exosomal subtype. Several studies have demonstrated that exosomes originating from immune cells possess immunostimulatory properties (Tang et al., 2016; Srinivasan et al., 2017) and can be utilized for immune activation and disease prevention. M1 macrophages not only present antigens to naive T cells but also secrete IL-12, thereby promoting the differentiation of T helper type 1 (Th1) immune responses. These cells play a crucial role in host defense against microbial pathogens and represent key components of the immune system (Philips and Ernst, 2012). In this study, the chicken macrophage cell line HD11 was stimulated with LPS, and qPCR was employed to assess the expression of typical M1 macrophage markers, including iNOS, IL-6, IL-12, IL-23, CD80, and CD86, which serve as established indicators for M1 phenotypic characterizatio (Atri et al., 2018; Yunna et al., 2020). This study demonstrated macrophage polarization to the M1 phenotype and successfully isolated exosomes from M1 chicken macrophages (HD11_M1_-EVs) using ultrahigh-speed centrifugation, with characterization confirming typical morphological and molecular profiles. Although previous research has established that exosomes derived from LPS-stimulated macrophages exhibit immunostimulatory effects at the cellular level (Hong et al., 2021a), their immunomodulatory potential at the organismal level remains unreported.

Phagocytosis represents a fundamental function of macrophages, and enhanced phagocytic capacity serves as a key indicator of macrophage activation status (Murray and Wynn, 2011). Our experimental results demonstrate that following 24-hour stimulation with HD11_M1_-exo, macrophages display significantly enhanced capacity to phagocytose neutral red, indicating that HD11_M1_-exo stimulates innate immune responses in macrophages. The Toll-like receptor (TLR) signaling pathway serves as a pivotal regulator of immune responses, operating through two distinct pathways: the MyD88-dependent pathway and the TRIF-dependent pathway (Duan et al., 2022). Established research has demonstrated that LPS stimulation of chicken macrophage-derived exosomes can induce immune responses and cytokine production through the MyD88/NF-κB signaling pathway (Hong et al., 2021a). However, the signaling mechanisms of M1 macrophage-derived exosomes via the TRIF pathway remain uncharacterized. Our experimental results demonstrate that HD11_M1_-exo activates the LPS/TLR4 signaling pathway by targeting upstream components of the TLR signaling cascade. HD11_M1_-exo mediates immunostimulatory functions through both MyD88-dependent and TRIF-dependent signaling pathways. This study elucidates novel regulatory mechanisms of avian M1 macrophage exosomes in the TLR signaling pathway.

To date, numerous studies have demonstrated successful enhancement of innate immunity in chicken embryos through administration of immunostimulants, which confer enhanced resistance against IBV infection. Bal Krishan Sharma et al. experimentally demonstrated the inhibitory effects of TLR agonists on IBV replication in chicken embryos, elucidating the underlying mechanism by which this protection is mediated through modulation of proinflammatory cytokine and antiviral gene expression (Sharma et al., 2020). TNF-α serves as a pleiotropic cytokine that mediates inflammatory responses, antitumor immunity, and immune homeostasis across diverse physiological processes (Aggarwal, 2003; Croft, 2009). The present study demonstrated that HD11_M1_-exo pretreatment (24 h prior to IBV challenge) significantly upregulated TNF-α gene transcription in embryonic tracheal tissues, enhanced immune competence, and mitigated IBV-induced pathological lesions. These findings indicate that HD11_M1_-exo induces immunostimulatory responses in avian embryos.

Furthermore, in vivo trials were conducted to evaluate HD11_M1_-exo immunomodulatory properties, which revealed that HD11_M1_-exo administration significantly upregulated CD80 and CD86 transcription in tracheal and Harderian gland tissues of vaccinated birds, thereby enhancing cell-mediated immune responses in avian tissues. As a critical component of the host defense network, the mucosal immune system serves a pivotal function in avian immunity. Birds employ their distinctive mucosal immune apparatus to mount effective immune responses against pathogen invasion, thus providing protection against infectious agents (Nochi et al., 2018). Immunoglobulin A (IgA), as the predominant antibody subtype in the mucosal immune system, is widely distributed on the mucosal surfaces of the gastrointestinal tract, respiratory tract, vagina, and in bodily fluids such as tears and saliva, forming a crucial defense line for the body’s mucosal immune protection (Li et al., 2020). For poultry, localized immunoglobulins in tears, such as IgA, are produced by lymphocytes in the Harderian gland, providing local protection to the upper respiratory tract tissues (Baba et al., 1988). Additionally, TGF-β4 can induce B cells to produce the non-inflammatory immunoglobulin subclasses IgG4 and IgA, playing a crucial role in IgA generation (Akdis et al., 2004). In our investigation, IgA concentrations in lacrimal fluid and TGF-β4 transcriptional activity in tracheal tissues of chickens exposed to HD11_M1_-exo were significantly elevated compared to control subjects. This study represents the first demonstration at the organismal level that avian M1 macrophage-derived exosomes activate both cell-mediated and mucosal immune responses in chickens, indicating that HD11_M1_-exo represents a novel candidate immunostimulant for veterinary applications.

The immunomodulatory effects of M1 macrophage-derived exosomes on vaccine-induced immunity represent a central research objective of this investigation. In this study, HD11_M1_-exo was evaluated as an adjuvant coadministered with the IBV live attenuated vaccine H120. Results demonstrated that HD11_M1_-exo/H120 coadministration significantly upregulated CD80 and CD86 transcription in tracheal and Harderian gland tissues at multiple time points post-vaccination. This combination also enhanced IgA concentrations in lacrimal fluid of vaccinated birds and increased TGF-β4 gene expression in tracheal tissues. Furthermore, H120-induced IgY antibody titers were augmented, with the HD11_M1_-exo group exhibiting significantly higher antibody levels at 14 days post-vaccination (dpv) compared to the adjuvant control group. These findings represent the first experimental evidence that HD11_M1_-exo coadministration with IBV live attenuated vaccine H120 significantly enhances cell-mediated, humoral, and mucosal immune responses to IBV vaccination, thereby potentiating vaccine efficacy. These data indicate that HD11_M1_-exo constitutes a novel vaccine adjuvant candidate for avian respiratory vaccines.

The focus of this study is to investigate whether HD11_M1_-exo can serve as an adjuvant to resist strong pathogenic infections of IBV and whether it can enhance the cross-protective effects of existing vaccines. Therefore, this study conducted animal experiments on the combination of HD11_M1_-exo and vaccines to resist IBV infection. The initial infection site of IBV is typically located in the upper respiratory tract, primarily targeting ciliated cells and mucous-secreting cells (Dhinakar Raj and Jones, 1996). Multiple studies have demonstrated that during IBV infection, the viral load in the trachea typically peaks at 5 days post-infection (**dpi**) and subsequently declines gradually (Okino et al., 2013). The IBV HX strain utilized in this study is a highly pathogenic strain isolated from Guangxi Province, China, exhibiting a 5% nucleotide sequence difference in the S1 gene, a threshold associated with insufficient cross-protection from commercial vaccines (Villarreal et al., 2010; Marandino et al., 2015). Our preliminary research demonstrates that sequence similarity analysis of the IBV HX strain with vaccine strains H120, 4/91, and LDT3-A reveals relatively low similarity of the S1 gene at both nucleotide (58.49%, 58.80%, and 59.65%, respectively) and amino acid (52.33%, 52.09%, and 53.51%, respectively) levels (Yang et al., 2024). Therefore, the HX strain provides an appropriate model to evaluate whether HD11_M1_-exo can enhance the cross-protective efficacy of the low-virulence vaccine H120. Our experimental results demonstrate that combining HD11_M1_-exo with the low-virulence IB vaccine H120 significantly reduces IBV viral load in the trachea of immunized chickens, significantly alleviating pathological damage in the trachea and mitigating clinical signs in infected chickens. This study provides the first evidence that exosomes derived from M1 macrophages can enhance the cross-protective efficacy of existing vaccines, effectively protecting against infections by virulent IBV strains. HD11_M1_-exo may serve as a vaccine adjuvant to provide protection for commercial poultry.

## Conclusion

In summary, this study investigates exosomes derived from LPS-stimulated M1 macrophages, providing the first comprehensive analysis at cellular, embryonated egg, and animal levels (Figure 14). Cellular-level analysis demonstrated that HD11_M1_-exo activates the LPS/TLR4 signaling pathway in macrophages, enhancing their immunological activity. In embryonated eggs, HD11_M1_-exo pretreatment increases TNF-α gene expression in the trachea, thereby enhancing resistance to IBV infection. At the animal level, HD11_M1_-exo significantly enhances cellular, humoral, and mucosal immunity in immunized chickens. Furthermore, challenge experiments demonstrated that HD11_M1_-exo, when used as a vaccine adjuvant, effectively enhances the cross-protective efficacy of existing IB vaccines against highly pathogenic IBV infections.

**Figure 14.**
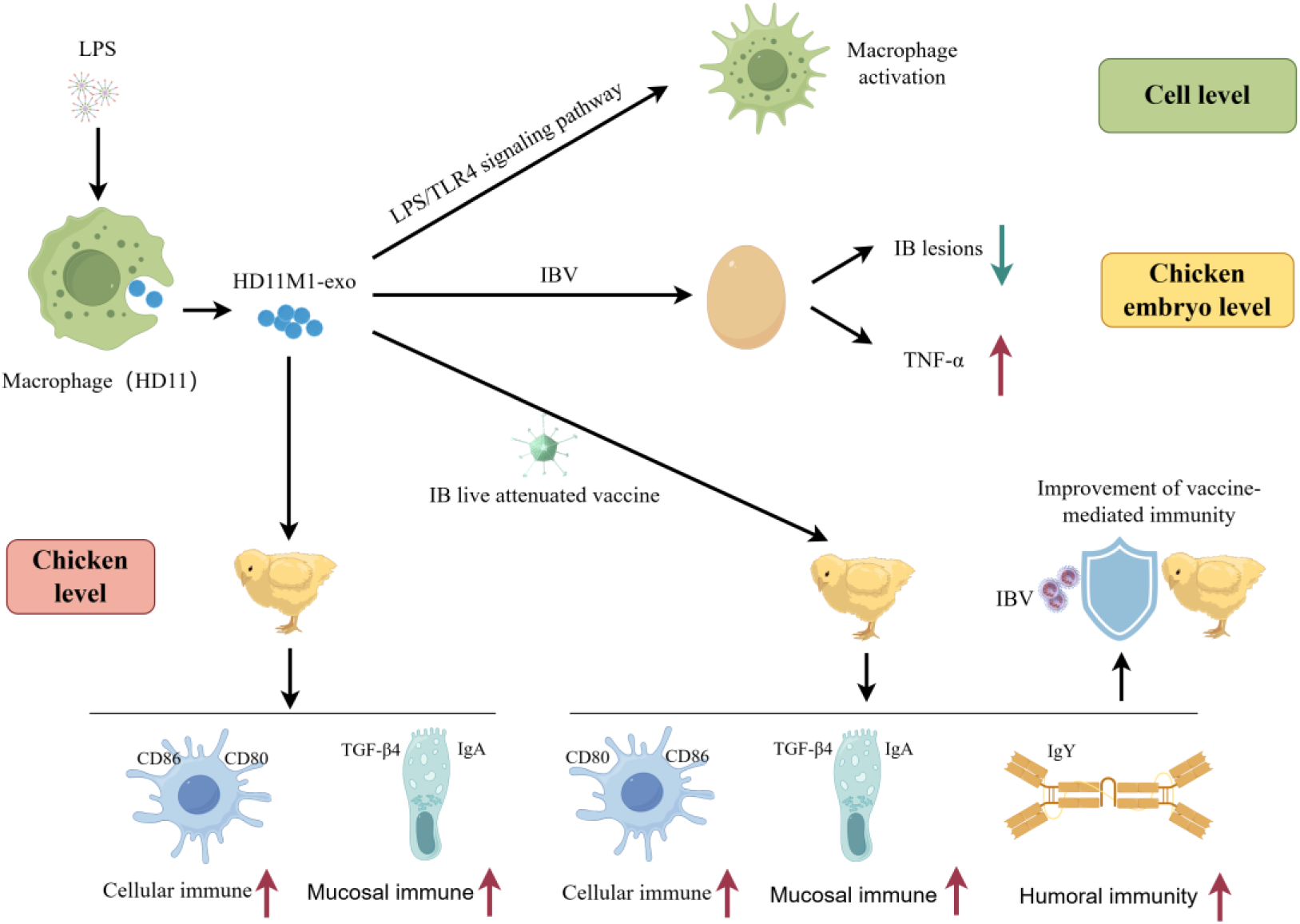
Schematic representation of the experimental design and workflow.

## Funding

The authors declare that financial support was received for the research, authorship, and/or publication of this article. This research was funded by National Natural Science Foundation of China (32202782), Guangdong Basic and Applied Basic Research Foundation (2021A1515012388).

## Conflict of Interest

The authors declare that the research was conducted in the absence of any commercial or financial relationships that could be construed as a potential conflict of interest.

